# Gene expression across tissues, sex, and life stages in the sea urchin *Tripneustes gratilla* [Toxopneustidae, Odontophora, Camarodonta]

**DOI:** 10.1101/284224

**Authors:** Áki Jarl Láruson, Simon E. Coppard, Melissa H. Pespeni, Floyd A. Reed

## Abstract

The pan-tropical sea urchin *Tripneustes gratilla* is an ecologically and economically important shallow water algal grazer. The aquaculture of *T. gratilla* has spurred growing interest in the population biology of the species, and by extension the generation of more molecular resources. To this purpose, *de novo* transcriptomes of *T. gratilla* were generated for two adults, a male and a female, as well as for a cohort of approximately 1,000 plutei larvae. Gene expression profiles of three adult tissue samples were quantified and compared. These samples were of gonadal tissue, the neural ring, and pooled tube feet and pedicellariae. Levels of shared and different gene expression between sexes, as well as across functional categories of interest, including the immune system, toxins, genes involved in fertilization, and sensory genes are highlighted. Differences in expression of Sex determining Region Y-related High Mobility Group box groups and general isoform expression between the sexes is observed. Additionally an expansion of the tumor suppressor DMBT1 was observed in *T. gratilla* when compared to the annotated genome of the sea urchin *Strongylocentrotus purpuratus*. The draft transcriptome of *T. gratilla* is presented here in order to facilitate more genomic level analysis of de-novo sea urchin systems.

## Introduction

The Phylum Echinodermata occupies a unique place on the evolutionary tree of life. Together the Echinodermata and Hemichordates, or acorn worms, form the clade Ambulacraria, which are the most closely related invertebrate taxa to the Chordates (Metschnikoff 1881, Furlong & Holland 2002, Satoh *et al*. 2014). Sea urchins (Class Echinoidea) have served as a model organism in developmental biology for over 150 years, owing to their frequent and voluminous broadcast spawning behavior and the ease with which gametes are collected and observed for *in vitro* fertilization (McClay 2011). The importance of sea urchins to the field of developmental biology resulted in an official call by Davidson & Cameron (2002) to sequence a complete sea urchin genome. In 2006 the full genome of the East Pacific urchin *Strongylocentrotus purpuratus* (Stimpson 1857) was published, and the annotated genome made freely available online (Sea Urchin Genome Sequencing Consortium 2006, Cameron *et al*. 2009). This development inspired significant and ongoing molecular work on members of the Strongylocentrotidae family (Kober & Bernardi 2013, Oliver *et al*. 2010, Walters *et al*. 2008), and more recently, a handful of transcriptomes and representative genomes for Echinoderms beyond the Strongylocentrotidae have been published (Dilly *et al*. 2015, Israel *et al*. 2016). While there is routine use of sea urchins as laboratory organisms and a growing body of molecular data available, many aspects of sea urchin biology still remain unknown. For example, while there have been hypotheses generated about the sex determination of sea urchins, it has yet to be concluded whether sea urchins operate under genetic sex determination or even possess distinct sex chromosomes (Eno *et al*. 2009; Bachtrog *et al*. 2014). Many species of sea urchin are also highly commercially valued in the food industry. Notably *Tripneustes gratilla* (Linnaeus 1758), commonly known as the collector urchin, and the Pacific congeneric of the Caribbean “sea egg” *T. ventricosus* (Lamark 1816). *Tripneustes gratilla* has one of the most expansive ranges of all shallow water echinoids, occurring everywhere from the Hawaiian islands in the central Pacific, to the shores of South Africa (Mayr 1954, but see Bronstein *et al*. 2017). As an ecosystem engineer, *T. gratilla* greatly affects the shallow water community composition during rapid population expansions and mass die-offs that frequently occur subsequent to rapid expansions (Valentine & Edgar 2010). As such, *T. gratilla* is currently the object of a large-scale aquaculture and outplanting effort by the state of Hawai◻i as a biocontrol agent against invasive algae such as *Acanthophoraspicifera, Gracilaria salicornia, Eucheuma denticulatum* and *Kappaphycus clade B* (Westbrook *et al*. 2014). A member of the family Toxopneustidae (Troschel 1872), a sister family to the Strongylocentrotidae, (Láruson 2017), *T. gratilla* is a confamilial of the highly venomous sea urchin, *Toxopneustes pileolus* (Lamark 1816). The increased collection and cultivation of *T. gratilla* has spawned increased interest in understanding the population structure of this broadly distributed urchin (Cyrus *et al*. 2014, Westbrook *et al*. 2015). This paper presents comparative descriptions of annotated draft transcriptomes from a male and female adult as well as a cohort of larval *T. gratilla* to determine how gene expression differs in both somatic and gonadal tissues between the sexes, and how gene expression profiles vary at different life stages (planktotrophic larvae versus benthic adults). A focus is placed on key similarities in presence or absence of expressed genes between these broadly split life stages. As suggested by Tu *et al*. 2012, the combination of larval developmental stage and adult tissue transcriptome can be integral for a more accurate context of genome level sequences. We present this draft transcriptome in response to the need for more large scale molecular resources for this ecologically and economically important sea urchin.

## Materials & Methods

### Collection and Extraction

Whole RNA was extracted using a Qiagen RNeasy extraction kit from two adult *T. gratilla*, collected from near-shore waters of southwest O◻ahu, Hawai◻i (approximately 21°21’13′N, 158°7’54′W). Three distinct tissues were individually sampled and indexed from each adult: gonadal tissue was sampled from each of the five gonadal lobes; neural tissue was sampled from the neural ring; tube feet and pedicellariae were sampled from several places across the animals’ external surface. Whole RNA was similarly extracted from approximately 1,000 plutei-stage *T. gratilla* larvae, acquired from the Ānuenue Fisheries Research Center on Sand Island, O◻ahu, Hawai◻i (Hawai◻i Department of Land and Natural Resources). The larval cohort consisted of mixed progeny from five adult females and five adult males. These parental *T. gratilla* were mature (>65mm in diameter) individuals wild caught off the coast of western O‘ahu, Hawai‘i.

### Sequencing

Extracted RNA was sent to the Hawai◻i Institute of Marine Biology Genetics Core for cDNA library construction and sequencing. Extracts were treated with Epicentre Rnase-free Dnase I; 1 uL of Dnase I was added per 20uL of sample, then incubated at 37°C for 30 min. Samples were then cleaned using the Qiagen RNeasy Minelute Cleanup Kit. Quality was assessed on an Agilent 2100 Bioanalyzer. Poly-A tail isolation and cDNA synthesis were performed with the Illumina TruSeq Stranded mRNA Sample Preparation Kit (Protocol Part # 15031047 Rev. E) with no fragmentation time to allow for larger RNA fragments. cDNAs were sequenced on two lanes of an Illumina MiSeq with single strand chemistry for 300 cycles, using a V2 reagent kit. Sequence files for each sample are available on the NCBI SRA database (Female Gonad: SRR6844874, Female Neural: SRR6844873, Female Tubefeet: SRR6844872, Male Gonad: SRR6844871, Male Neural: SRR6844877, Male Tubefeet: SRR6844876, Larvae: SRR6844875).

### Sequence Assembly, Annotation, and Analysis

All sequence reads were filtered and trimmed using Trimmomatic (Bolger et al. 2014): matches to TruSeq3 index adapters were purged, leading and trailing bases falling under a Phred33 quality score of 3 were trimmed, a sliding window trim was performed with a window size of four bases and an average window quality score threshold of 15, and a minimum read threshold of 36 was set. Cleaned reads from each sample were then assembled individually with Trinity (Haas *et al*. 2013). In order to avoid an excess of falsely identified isoform variants resultant assembly was clustered at a 99% identity threshold with CD-HIT (Fu *et al*. 2012, Li & Godzik 2006). Assembly quality was partly assessed via recovery of conserved metazoan single copy orthologs, as defined through the software BUSCO (Simão *et al*. 2015). Reciprocal best matches to the annotated genomes of *S. purpuratus* (Spur4.2) and *Lytechinus variegatus* (Lvar2.2), acquired from www.echinobase.org, were performed with a local tBLASTx alignment with an e-value cut off of 1e-10. Additional annotation was performed by mapping the assembly, as well as the longest open reading frames from Transdecoder (Haas *et al*. 2013), to the SwissProt (Bairoch & Apweiler 2000) and Pfam (Finn *et al*. 2016) databases. The resulting hits were assembled into a database with Trinnotate (https://trinotate.github.io/), with Gene Ontology (GO) categories assigned to said hits. Transcript quantification, in terms of Transcripts Per kilobase Million (TPM), was accomplished with RSEM (Li & Dewey 2011). Manhattan distance values, summary statistics, and visualization was generated in R (R Core Team 2017)..

## Results

In total, 14,903,815 raw sequence 300bp reads were generated. Trinity assembly of sequence data clustered at 99% identity from all seven samples resulted in 172,841 transcripts, with 157,610 flagged as unigenes. The final assembly had an average contig length of 721.09, an N50 of 917, and a GC content of 37.04%. Assembly quality was partly assessed via the recovery of 86.6% of highly conserved metazoan single copy orthologs. For comparison, the full *S. purpuratus* genome recovered 90.3% of the same dataset of metazoan orthologs. Only 10,027 unigenes had proposed Trinity identified splice variants, or isoforms. Some 11,689 unigenes were annotated via reciprocal best matches to *S. purpuratus*, while an additional 3,519 unigenes were annotated via matches to *L. variegatus* for a total of 15,208 annotated unigenes. Of the remaining 142,402 unannotated unigenes, 6,452 matches to the SwissProt and Pfam databases for a total of 21,660 annotated transcripts. This corresponds to just over 70% (21,660/29,948) of the total number of annotated genes of the *S. purpuratus* genome.

Between all annotated genes over 56% were observed exclusively in the adult tissues, and only 2% (422 genes) were uniquely observed in the larval transcriptome (Figure 1A). 86.8% of all annotated genes were observed in both individuals, with 6.9% of annotated genes being unique to the female, and 6.3% being unique to the male (Figure 1B). 58% of all annotated genes were found to be co-occurring across the three adult tissues (gonad, neural, and tube feet/pedicellariae). The largest number of tissue-specific expression was seen in the gonads (7.7%), while the fewest uniquely expressed genes were identified in the tube feet/pedicellariae (1.3%, Figure 1C).

**Figure 1.**
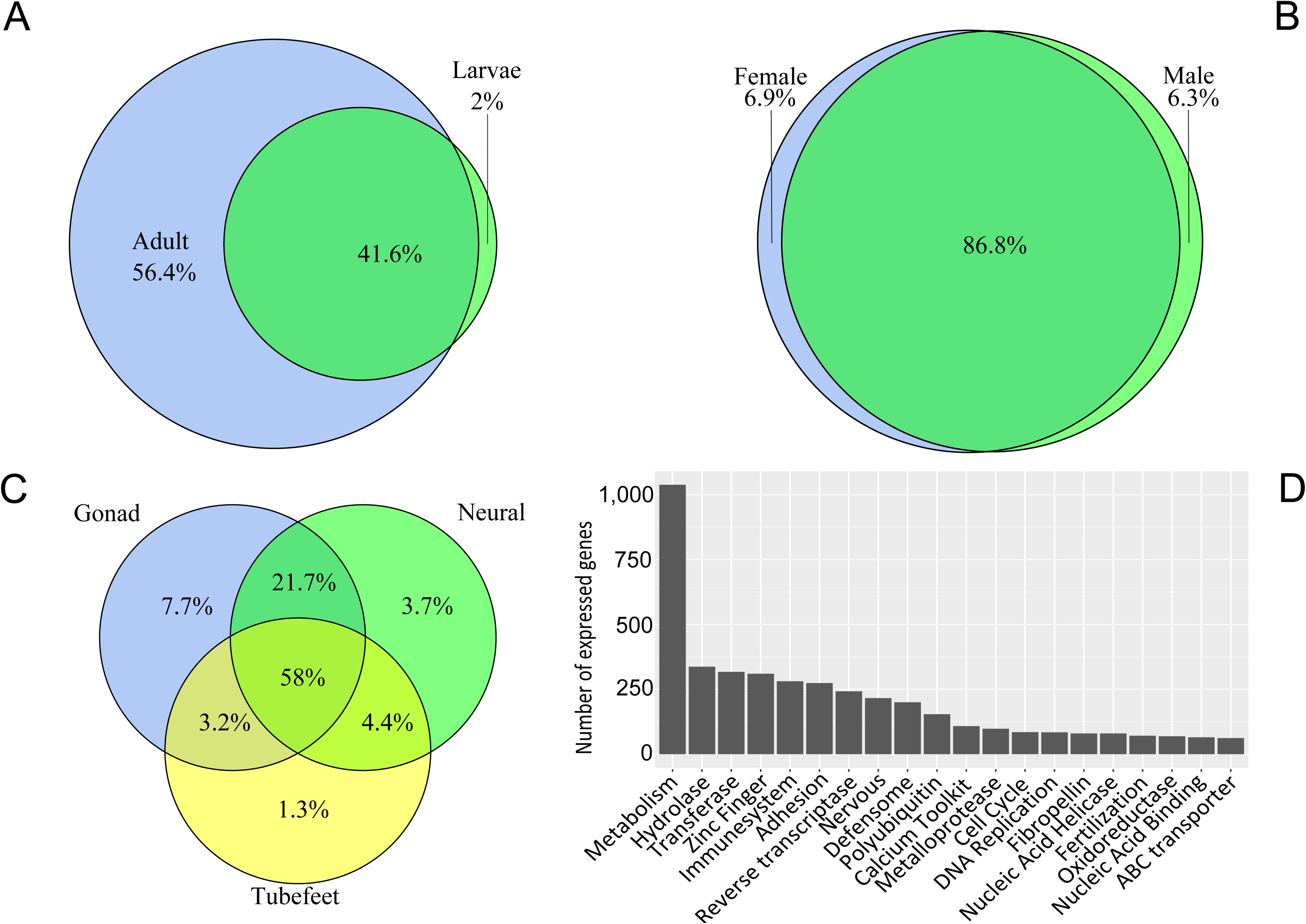
Annotated genes across samples. A. Genes identified across adult tissues versus larval sample. B. Genes identified across all tissues in a female versus in a male. C. Genes identified across different tissues. D. Number of identified genes expressed across functional categories identified via annotation to *Strongylocentrotus purpuratus, Lytechinus variegatus*, as well as Pfam and Swissprot datasets. Only the top 20 largest categories are displayed.

Across broadly identified functional categories, after excluding ribosomal genes as well as genes with unknown annotations, by far the greatest number of annotated genes were identified as being involved in metabolism (1,038). The 20 most highly represented functional gene categories are highlighted in Figure 1D.

### Fertilization and Sex differences

A Manhattan distance calculation of expression profiles between adult tissues clusters each adult separately, with the neural and tube feet/pedicellariae tissues clustering more closely to each other than to the gonadal tissue. However when isoforms are clustered so that only unigene expression information is considered, the variation between individual gonad expression profiles is reduced such that the male and female gonad samples cluster together (Figure 2). All annotations and expression profiles across tissues are made available in Supplementary Table 1.

**Figure 2.**
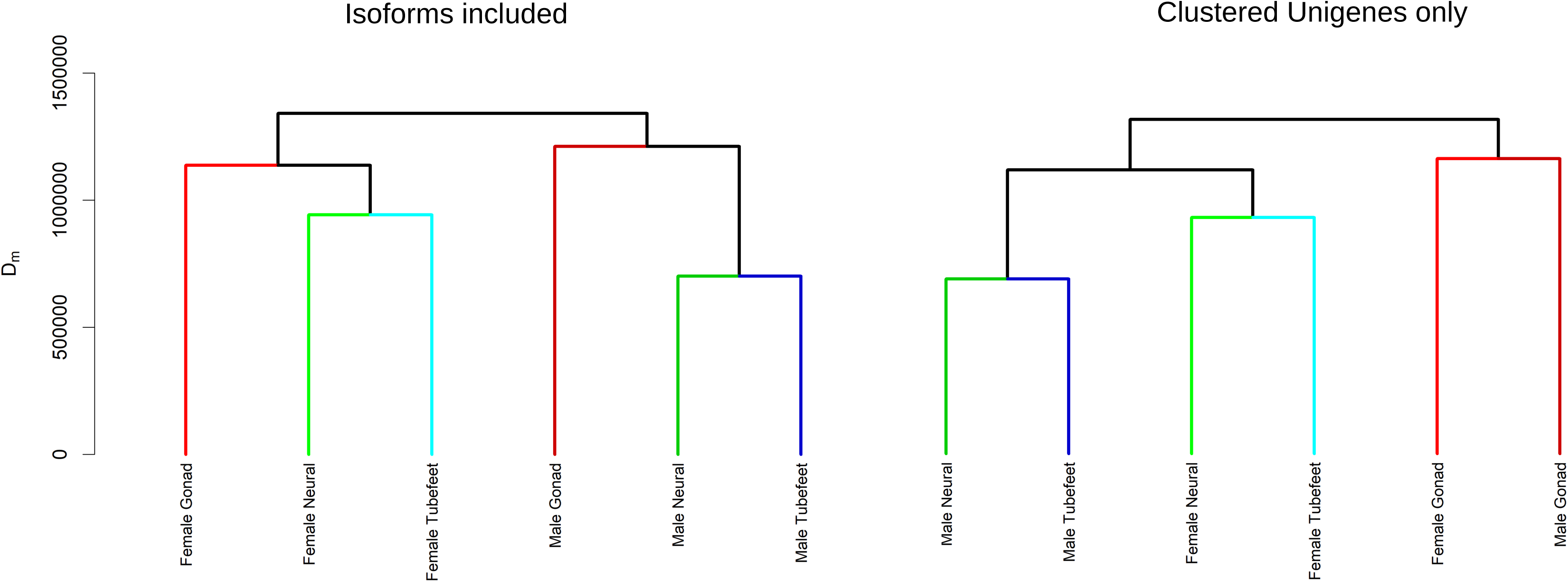
Dendrogram of Manhattan distances observed between expression profiles in adult tissues. On the left including all isoforms identified in each sample, and on the right considering only the expression of unigenes (clustered isoforms). The placement of Female Gonad and Male Gonad profiles as a distinct unit when considering only the unigene expression suggest that splice variation underlies the differences in male and female gonadal expression.

Ninety genes were identified as being involved in fertilization, with 42 of those expressed in both the adult samples as well as the larvae. Two genes, acid-sensing ion channel-like (AsicL) and sodium channel (NaC) were only observed in the larvae. The dual oxidase homolog Udx1, which has an important role in the fast-block to polyspermy (Wong *et al*. 2004), was one of the four most highly expressed fertilization genes across all samples, except for in the male gonad. Comparing male and female samples across all tissues recovered 1,335 genes that were exclusively found in the male and not the female, and 1,467 genes were only seen in the female and not the male. The only sex determination associated protein recovered in one sex was Wnt-4, which was found in the male and the larvae. Eleven Sex determining Region Y related High Mobility Group box (Sox) gene fragments were recovered. These represented the SoxB1, SoxB2, SoxC, SoxD, SoxE, SoxF, and SoxH groups; these groups were seen across all adult tissues, while SoxD was the only group not recovered from the larvae (Figure 3).

**Figure 3.**
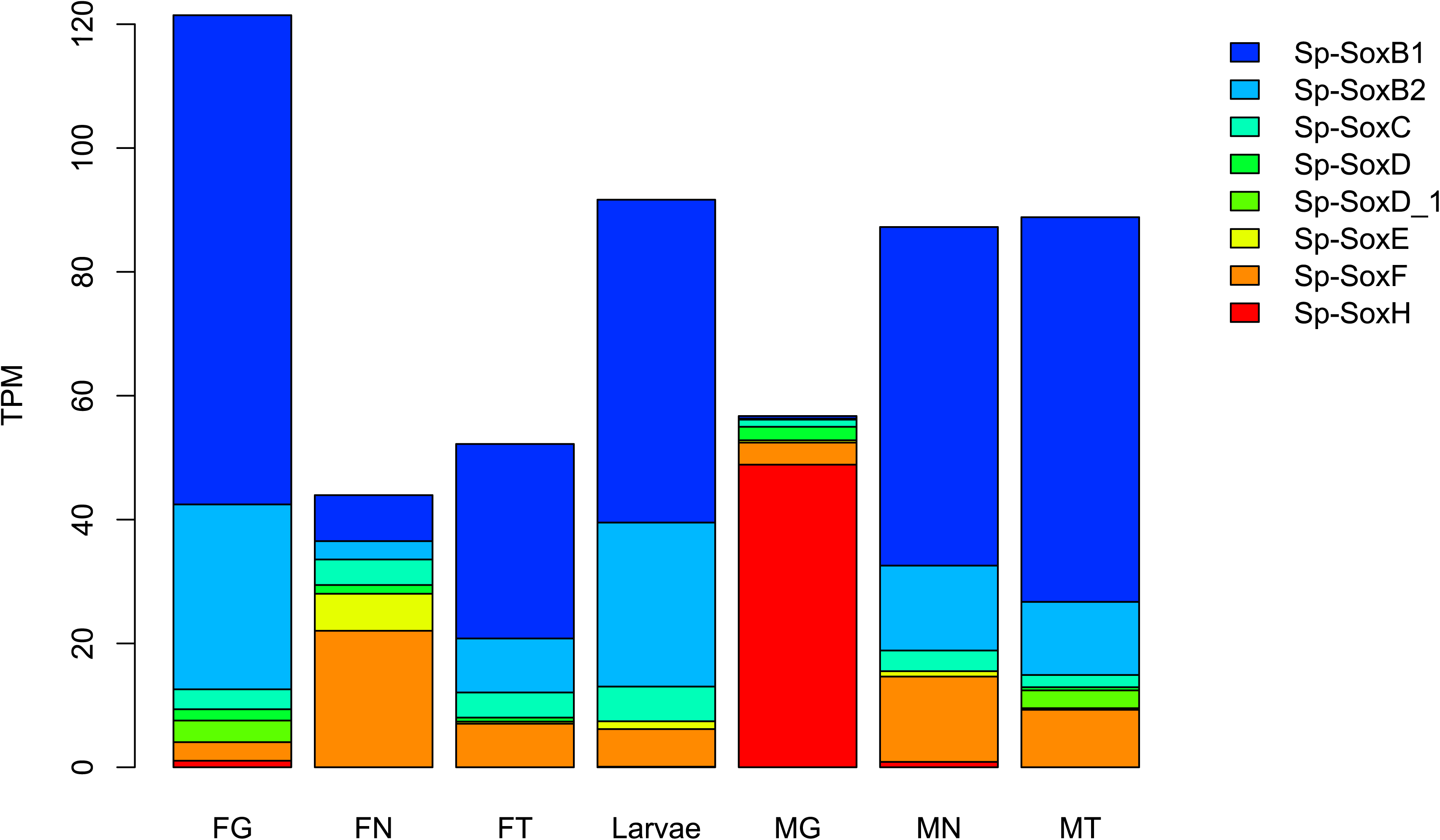
Barplot of Transcripts Per kilobase Million levels of Sex determining Region Y-related High Mobility Group box (SRY-HMG or SOX) group expression across the different samples. Elevated SoxH expression was observed in the Male Gonad compared to the remaining samples, while higher levels of SoxB1, SoxB2, and SoxC were observed in the female and the larvae. F = Female, M=Male, G=Gonad, N=Neural, T=Tubefeet/Pedicellariae.

### Immune system

Immune system genes were identified via known annotated functional category (Hibino *et al*. 2006) or GO categorization. Comparing only the most highly expressed immune genes, Ubiquitin-40S ribosomal protein S27a (RPS27A) fusion protein was the most highly expressed in all female tissue samples, the larvae sample, as well as the male neural and tube feet/pedicellariae samples. In the male gonad sample, however, a Scavenger Receptor Cysteine Rich (SRCR) gene (Srcr197) was the most highly expressed immune gene observed. SRCR domains were seen expressed in all samples, with 33 unique SRCR genes identified in total. Two genes involved in the clotting response, Amassin (Hillier & Vacquier 2003), and a Kringle domain containing gene (PlgL2), were observed to be two of most highly expressed genes in the neural tissue of both the female and male samples. Additionally, PlgL2 was observed to be present at elevated levels in the tubefeet/pedicellariae from both adult samples. In total, nine sea urchin TLR genes were identified, along with six Tlr-like genes. Tlr003 was identified through the SwissProt database, but failed as a reciprocal best match to that of either the *S. purpuratus* or *L. variegatus* genomes.

Twenty-nine NACHT domain and leucine-rich repeat (NLR) proteins were identified. Overall 15 elements annotated to the immunoglobulin superfamily (IgSF) and 12 genes connected to V(D)J recombination were identified, as well as 18 tumor necrosis factor (TNF) functional category genes. Two RAS domains and two Fat-like cadherin-related tumor suppressor homologs were also identified. An unexpected finding was the identification of 69 Deleted in Malignant Brain Tumor 1 (DMBT1) unigenes. The majority of expressed DMBT1 genes were observed only in the adult tissues, but eight were also identified in the larvae. Results are summarized in Table 1.

**Table 1.** Identified unigenes from select functional gene classes implicated in immune system response.

| Gene class | Number of unigenes identified |
| --- | --- |
| Deleted in malignant brain tumors 1 (DMBT1) | 69 |
| Scavenger receptor cysteine-rich repeat (SRCR) | 33 |
| NACHT domain and leucine-rich repeat (NLR) | 29 |
| Tumor necrosis factor (TNF) | 18 |
| Toll-like receptor (TLR) | 15 |
| Immunoglobulin superfamily (IgSF) | 15 |

### Toxins

Six gene fragments annotated as putative toxins were identified through the Swissprot/Pfam database. Venom Protein 302 was identified in all adult tissues, absent in the larvae, but expressed most highly in the tube feet and pedicellariae samples of both adults. Snake venom metalloprotease inhibitor 02A10 was identified in the male gonad and Venom serine protease Bi-VSP was found in all adult tissues except for the male gonad. C-type lectin lectoxin-Enh5 was found in all female tissues, the larvae, and the male neural tissue. Fourteen transcripts with Phospholipase A2 (PLA2) activity that were most highly represented in the tubefeet/pedicellariae were identifed. Fragments annotated as Stonustoxin alpha subunit and Verrucotoxin beta subunit, both toxic proteins characterized from Stonefish (genus *Synanceia*), were expressed in the female tube feet/pedicellariae sample (Table 2). Alpha-latrotoxin-Lhe1a was also found in the female tube feet/pedicellariae. This fragment contained ankyrin repeats typically seen in Alpha-latrotoxin-Lhe1, but was missing the N-terminal domain associated with the neurotoxin from the western black widow spider *Latrodectus hesperus* Chamberlin & Ivie, 1934.

**Table 2.** Venom/Toxin associated gene fragments identified. Venom Protein 302 was the only transcript observed at its highest expression in the tube feet/pedicellaria of both individuals. All values reported in Transcripts Per kilobase Million.

| Transcripts | Annotation | F. Gonad | F. Neural | F. Tubefeet | Larvae | M. Gonad | M. Neural | M. Tubefeet |
| --- | --- | --- | --- | --- | --- | --- | --- | --- |
| TR37298_c0_g1 | Venom protein 302 | 0.67 | 5.2 | 15.27 | 0 | 0 | 7.54 | 10 |
| TR47326_c0_g2 | Stonustoxin subunit alpha | 0 | 0 | 18.3 | 0 | 2.07 | 0 | 0 |
| TR47326_c0_g3 | Stonustoxin subunit alpha | 0.52 | 0 | 6.61 | 0 | 0 | 0 | 0 |
| TR47326_c1_g1 | Verrucotoxin subunit beta | 0 | 0 | 10.92 | 0 | 0 | 0 | 0 |
| TR9549_c0_g1 | Alpha-latrotoxin-Lhe1a | 0 | 0 | 7.02 | 0 | 0 | 0 | 0 |
| TR80588_c0_g1 | Venom serine protease Bi-VSP | 6.68 | 4.68 | 4.22 | 0 | 0 | 10.93 | 5.5 |
| TR48480_c0_g1 | C-type lectin lectoxin-Enh5 | 2.32 | 0.9 | 0.81 | 2.21 | 0 | 2.81 | 0 |
| TR8624_c1_g2 | Snake venom metalloprotease inhibitor 02A10 | 0 | 0 | 0 | 0 | 31.3 | 0 | 0 |

### Sensory

Only four of the eight prototypical sea urchin opsin genes (Raible *et al*. 2006, D’Aniello *et al*. 2015, Lowe *et al*. 2017) were recovered: opsin 2, opsin 5, opsin 3.2, and opsin 4-like-1. All opsins were only found expressed in adult tissues. The most highly expressed sensory protein identified in the larvae was mechanosensory protein 2, which was also the most highly expressed sensory protein in the male and female tube feet/pedicellariae as well as the male neural tissue. Six distinct Sensory GPCR Rhodopsins and a FAD binding blue light photoreceptor sequence were recovered from adult as well as larval sequences.

## Discussion

Some 21,660 unigenes identified from a larval cohort, as well as three tissues of an adult male and female, are presented here from *Tripneustes gratilla*. This represents over 70% (21,660/29,948) of the total number of annotated genes of the *S. purpuratus* genome, and includes 86.6% of highly conserved metazoan single copy orthologs (Simão *et al*. 2015). This represents a significant portion of the *T. gratilla* transcriptome.

Since the sex determination mechanism of sea urchins remaining unclear, expression profile differences between three individually extracted tissue samples from a male and female *Tripneustes gratilla* provides some insight into potentially contributing molecular factors. The distinct gene expression profiles of the male and female gonads suggested by the Manhattan distance on clustered unigenes, disappears when measuring differences in expression profiles considering isoforms (Figure 2). While differences were observed between isoform expression profiles in male and female tissues, no isoforms were found to exclusively occur in one sex. This suggests that similar genes are expressed in both the male and female gonads, and levels of expression of splice variants are primarily contributing to the functional distinction of testis and ovary. While the major sex determination protein SoxA (Sex determining Region Y) is absent in the presented *T. gratilla* dataset, as well as the *S. purpuratus* and *L. variegatus* transcriptomes (Cameron *et al*. 2009), seven of the remaining eight Sox gene groups are represented in this dataset (SoxB1, SoxB2, SoxC, SoxD, SoxE, SoxF, and SoxH). Somewhat curiously, the expression patterns of Sox gene groups in the larvae much more closely resembled that of the female than that of the male tissues (Figure 3). Out of all Sox gene expressions, SoxH was found to be most highly expressed in the male compared to the female, with 48.87 TPM observed in the male gonad, compared to 1.06 TPM in the female gonad. A homolog to the human Sox30, SoxH has been suggested to be involved in the differentiation of male germ cells (Osaki *et al*.1999). Wnt-4, generally understood in mammals to be involved in the suppression of masculinization in early development (Chassot *et al*. 2012), was recovered in the larvae as well as in adult male neural tissue. It was however not found to in the adult female. It is therefore possible the functional distinction of Wnt-4 may have the opposite effect, promotion of masculinization, in sea urchins.

The small relative number of uniquely larval expressed genes could be a result of the expression profile of several adult tissues being compared to a single developmental stage larvae; alternatively it could possible be an effect of sequencing effort: while there was effectively equal sequencing effort in this study for the larval and the adult tissue sequencing (one illumina MiSeq lane each), a comparison to *S. purpuratus* datasets with seven fold greater sequencing effort for the larvae (2 lanes for adult, 14 for larvae, Pespeni et al. 2013a, Pespeni et al. 2013b) showed virtually no uniquely adult expressed genes, and over 32% of annotated genes being observed only in the larvae (supplementary figure 1).

This annotated draft transcriptome of *T. gratilla* captured some key immune system genes. Interestingly, only 15 sea urchin TLR and TLR-like genes were identified across all samples. This is a low number compared to the 222 TLR genes identified in *S. purpuratus* (Sea Urchin Genome Sequencing Consortium 2006), and more in line with the 10 genes observed in *Homo sapiens* Linnaeus, 1758 or the 28 *S. purpuratus* TLR transcripts recovered by Tu *et al*. 2012. Similarly, 33 unique SRCR domains identified in total, across all samples, which is a greater number of SRCR genes than the 16 genes in *H. sapiens*, but still unlikely to represent the full suite of SRCR diversity, as 218 SRCR genes have been identified in *S. purpuratus* (Sea Urchin Genome Sequencing Consortium 2006). However, an expansion was observed in a protein member of the SRCR superfamily: deleted in malignant brain tumor 1 (DMBT1), which was represented by 69 unigenes. Only 1 DMBT1 and 11 DMBT1-like genes have been identified in the *S. purpuratus* genome.. DMBT1 are known to be involved in regulating both the immune system and tumor cells (via apoptosis or immune response to tumor cells). Tumor suppressing proteins such as Tumor Necrosis Factor (TNF) and Rassf5 were also observed in all adult tissues as well the larvae. This enhanced diversification of immune and tumor suppressing genes may be involved in the suppression of transmissible cancer cells in a marine environment (Metzger *et al*. 2016). Furthermore, this could suggest alternative immune response strategies between *S. pupuratus* and *T. gratilla*: amplification of TLR genes (but see also Tu et al. 2012) versus DMBT1 duplication. Additionally, the role of immune system genes in sea urchin tissue regeneration is worthy of future study (Ramírez-Gómez *et al*. 2008).

A number of potentially toxic proteins were identified in this dataset. Being a member of family Toxopneustidae, *T. gratilla* is closely related to the most venomous sea urchin *Toxopneustes pileolus*. Venom isolated from the globiferous pedicellaria of *T. gratilla* has been shown to be lethal to mice and rabbit subjects, with respiratory distress and terminal tonic convulsions reported prior to death (Alender 1963). *Tripneustes gratilla* is not considered hazardous to humans, however irritation of the skin after handling adult individuals has been observed by the authors. Also highly expressed in the tubefeet and pedicellariae were fourteen PLA2 activity genes. PLA2 share structural similarity with the *T. pileolus* Contractin A venom (Hatakeyama *et al*. 2015). Venom Protein 302, an insulin-like growth factor binding protein, originally identified in the venom gland of the Chinese Swimming Scorpion (*Lychas mucronatus* (Fabricus 1798) was observed in all tissue samples, aside from the male gonad and the larvae. It is the only annotated toxin to be present at its highest expression value in the tube feet/pedicellariae of both adult individuals.

Verrucotoxin is a dimeric lethal toxin identified in the Stonefish *Synanceia verrucosa* (Garnier *et al*. 1995). Verrucotoxin subunit B annotated fragments were observed exclusively expressed in the tube feet/pedicellaria; however only in the female adult. Two transcript variants of another Stonefish toxin, Stonustoxin subunit B (Poh *et al*. 1992), were also observed in the female adult’s tube feet/pedicellariae. While Stonustoxin has previously been considered a novel marine toxin (Ghadessy *et al*. 1996), more recent analysis suggests that it may in fact have more broad functional ancestry, and as is true for many toxins need not necessarily have a toxic function across broader taxonomic identification (Ellisdon *et al*. 2015). The expression location of these supposed toxins in the tube feet/pedicellariae could suggest the underlying function of toxic defense against predation, the globiferous pedicellaria of *T. gratilla* are defensive structures (see review in Coppard *et al*. 2012) and are potentially pursuit-deterrents to predatory fish (Sheppard-Brennand, *et al*. 2017). Future efforts will be needed to fully isolate and confirm the effects of these potential toxins.

Lastly it should be mentioned that a large number of reverse transcriptase genes (242) were identified in the dataset (Figure 1D). Sea Urchin Retroviral-Like (SURL, Mag family of Ty3/Gypsy retrotransposons) mRNA recovered suggesting that this TE is still quite active in *T. gratilla*. (Springer *et al*.1991, Gonzales and Lessios 1999)

## Conclusion

The contribution of this *T. gratilla* transcriptome, with 21,660 identified unigene sequences, will allow for further research into comparative genetic characterization of echinoids, as well as molecular diversity and population structure of this broadly distributed, economically, and environmentally important species.

## Data repository

All raw sequence reads are to be deposited to the NCBI SRA database (https://www.ncbi.nlm.nih.gov/sra/SRP135810). The full table of gene fragments, annotations, and corresponding expression values is available as Supplementary Table 1.

## Acknowledgements

This research was funded and made possible by the Jessie D. Kay Memorial Fellowship; the Elizabeth A. Kay Endowed Award; the Charles H. & Margaret B. Edmondson Research Fund; the Watson T. Yoshimoto Fellowship, and the Hampton & Meredith Carson Fellowship, both administered by the Ecology, Evolution, and Conservation Biology specialization program at the University of Hawai◻i at Mānoa. Thanks to Victoria Sindorf, for helpful comments and suggestions. Special thanks to David Cohen and staff of the Ānuenue Fisheries Research Center (DLNR), and to Amy Eggers, of the Hawai◻i Institute of Marine Biology Genetics Core.

---

Supplementary figure 1. S. purpuratus dataset with seven fold greater sequencing effort for the larvae (2 lanes for adult, 14 for larvae) to compare with Fig. 1A.

