## Supplementary figures and images for "Gene expression across tissues, sex, and life stages in the sea urchin *Tripneustes gratilla* [Toxopneustidae, Odontophora, Camarodonta]"

### Supplementary Materials

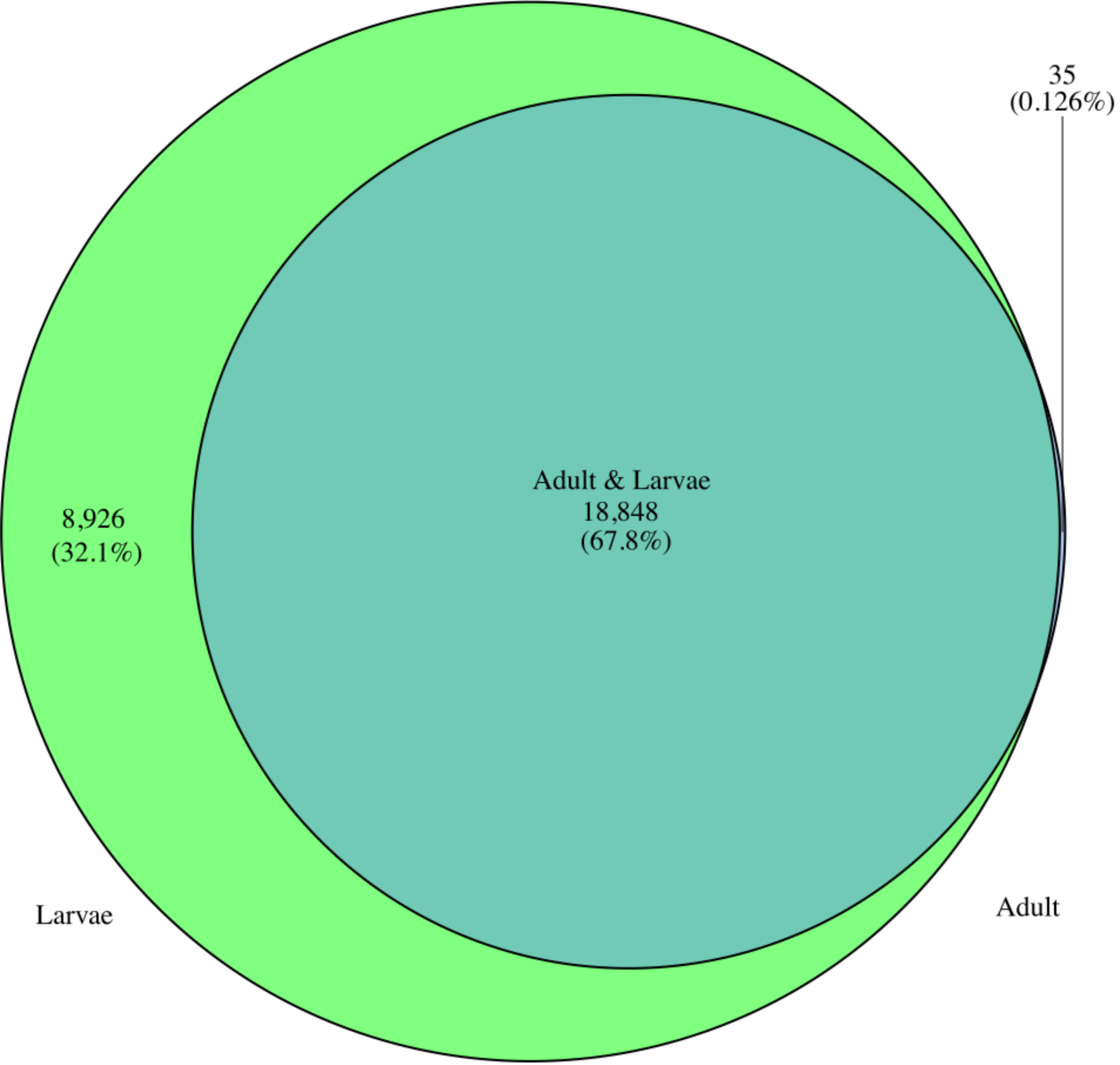
